# Categorization and discrimination of human and non-human primate affective vocalizations: investigation of the frontal cortex activity through fNIRS

**DOI:** 10.1101/2022.01.29.478308

**Authors:** C. Debracque, L. Ceravolo, Z. Clay, D. Grandjean, T. Gruber

## Abstract

Many species, including humans and non-human primates, react differently to threatening or pleasant situations. Because of its adaptiveness, recognizing affective signals is likely to be reflected in a capability of modern humans to recognize other closely related species’ call content. However, at both behavioural and neural levels, only few studies have used a comparative approach to understand affective decoding processes in humans, particularly with respect to affective vocalizations. Previous research in neuroscience about the recognition of human affective vocalizations has shown the critical involvement of temporal and frontal regions. In particular, frontal regions have been reported as crucial in the explicit decoding of vocal emotions especially in different task complexity such as discrimination or categorization. The aim of this study using functional Near Infrared Spectroscopy (fNIRS) was to specifically investigate the neural activity of the inferior frontal cortex *pars triangularis* (IFG_tri_) and the prefrontal cortex (PFC) underlying categorization (A versus B) and discrimination (A versus non-A) mechanisms of positive and negative affects in human, great apes (chimpanzee and bonobo), and monkey (rhesus macaque) vocalizations. We also analysed participants’ behavioural responses and correlated them with the recorded frontal activations. While performing the tasks, fNIRS data revealed a clear distinction between the two frontal regions, with a general positive activation of IFG_tri_ compared to a decrease of PFC activity. We also found a modulation of IFG_tri_ and PFC activations depending on both the species considered and on task complexity; with generally more activity in the IFG_tri_ during discrimination compared to categorization, and a more intense decrease of the PFC in categorization compared to discrimination. Behaviourally, participants recognized almost all affective cues in all species vocalizations at above chance levels in the discrimination task (except for threatening bonobo calls). For categorization, they mostly correctly identified at levels significantly above chance affective contents in human and great ape vocalizations but not in macaque calls. Overall, these findings support the hypothesis of a pre-human origin of affective recognition processing inherited from our common ancestor with other great apes and processed in the frontal cortex. Our results also highlight behavioural differences related to task complexity, i.e. between categorization and discrimination processes, and the differential involvement of the PFC and the IFG_tri_, which seems necessary to explicitly decode affects in all primate vocalizations.

## Introduction

Human life is made of choices, especially in the social domain. How we should react to threatening or joyful voices expressed by others conditions how we thrive in a given society. While usually associated with irrational choices, emotions are in fact essential to guide cognitive processes to enable adaptive responses to the environment (Brosch et al., 2013). Over the last three decades, researchers in psychology (for a review, see Lerner, Li, Valdesolo, & Kassam, 2015) and neurosciences (for a review, see Phelps, Lempert, & Sokol-Hessner, 2014) have investigated the impact of emotions on decision-making processes. Far from being only limited to humans, there is also a deep evolutionary origin to such recognition mechanisms. Allowing animal species to evaluate social motivations of others (Albuquerque et al., 2016) and then to react adaptively to a pleasant or a dangerous situation (Mendl & Paul, 2020), these recognition mechanisms are crucial for the fitness of individuals (Anderson & Adolphs, 2014; Filippi et al., 2017). In fact, perhaps even more importantly than for our own species (*Homo sapiens*), to correctly identify an affective signal in vocalizations is often a matter of life or death in the animal kingdom. For example, research on non-human primates (from henceforth, primates), our closest relatives, have demonstrated the capacity of chimpanzees to distinguish between different kinds of calls as function of the severities of aggression (Slocombe et al., 2009). Similar results have been found in other primates, with Gouzoules reporting the abilities of macaques to differentiate the seriousness of an agonistic interaction while listening to the victim’s calls (Gouzoules, 1984).

Recent research in humans on these recognition mechanisms has emphasized the role of available sensory information as well as the different levels of complexity involved in the process during which a human makes a decision among several options (de Lange & Fritsche, 2017). In particular, perceptual decision-making involves processing sensory information, which are evaluated and integrated according to the goal and the internal state of an individual but also depending on the possible number of choices (Hauser & Salinas, 2014). An important aspect of this research is to investigate the cerebral basis of such recognition. However, neuroscience studies have mainly focused on the visual domain. Therefore, the neural bases of perceptual decision-making using affective auditory information remain to be investigated.

Until now, functional Magnetic Resonance Imaging (fMRI) studies involving explicit recognition of affective cues in voices have emphasized the role of frontal regions, such as the inferior frontal cortex (IFG). For instance, Brück and colleagues have revealed a stronger activation in the IFG when the participants were explicitly decoding emotional prosody as compared to identifying phonetic or semantic aspects of speech (Brück et al., 2011). These results are in line with previous research showing a key role of the IFG in affective prosody decoding (Ethofer et al., 2006; Wildgruber et al., 2009). Furthermore, recent findings have highlighted the role of the IFG in the complexity of perceptual decision-making. The categorization (unbiased choice, ‘A vs B’) or the discrimination (biased choice, ‘A vs non-A’) of affective cues in voices indeed involves different subparts of the IFG, with the involvement of the *pars triangularis* (IFG_tri_) for discrimination and the involvement of the *pars opercularis* (IFG_oper)_ for categorization respectively (Dricu et al., 2017).

Unlike IFG, the role of the prefrontal cortex (PFC), well-known for its involvement in decision-making (e.g. Brosch et al., 2013; Damasio, 1996), remains poorly explored in regards to the vocal decoding of emotions. Yet, the emergence of functional Near Infrared Spectroscopy (fNIRS), a non-invasive technique to study the brain hemodynamic (Boas et al., 2014) using the principle of tissue transillumination (Bright, 1831), may shed new lights on these processes. Indeed, fNIRS studies have investigated the role of PFC in emotional processing, highlighting its role in emotion regulation (Glotzbach et al., 2011) and emotion induction (Matsuo et al., 2003; Ohtani et al., 2005; Yang et al., 2007). Interestingly, recent fNIRS studies pointed out the roles of both PFC and IFG in the vocal decoding of emotions. For instance, Zhang and colleagues reported a strong involvement of the human PFC and IFG during the discrimination of affective voices (Zhang et al., 2018). Similarly, Gruber and colleagues highlighted the modulation of IFG activity depending on the categorization or the discrimination of affects in auditory stimuli (Gruber et al., 2020). Hence, more investigations on PFC and IFG activations are necessary to improve our knowledge of affective decoding. Moreover, the fNIRS methodology seems particularly adapted to the exploration of frontal regions in decision-making and emotional paradigms.

Interestingly, anatomical structures (Petrides & Pandya, 2002; Rolls, 2004) and functions of the IFG and PFC in decision-making, auditory and affective processing are shared by most primate species, (e.g. macaques - *Macaca mulatta*; see Barbas, 2000; Barbas et al., 2011; Binder et al., 2004; Davidson, 1992; Frühholz & Grandjean, 2013; Kambara et al., 2018; LeDoux, 2012). In addition, as members of the *Hominidae* clade, which appeared between 13 and 18 million years ago (Perelman et al., 2011), modern humans share with the other living great apes (chimpanzees - *Pan troglodytes*, bonobos - *Pan Paniscus*, gorillas - *Gorilla subs*, and orangutans - *Pongo subs*) a large frontal cortex (Semendeferi et al., 2002). Overall, the fact that both humans and non-human primate species are able to identify correctly affective cues in conspecific vocalizations allowing them to use available information to make their choices; and that there is an anatomic and potentially functional convergence of the IFG and PFC across primate species, suggest that a comparative approach is particularly of interest to investigate the current role of these frontal regions in the human recognition of vocal emotions. Such approach may rely on primate calls beyond human vocalizations to uncover the evolutionary of human evaluation processes.

Yet, only a few studies have used a comparative approach to understand affective decoding mechanisms in humans using primate vocalizations. These studies have revealed at both cerebral and behavioural levels promising results highlighting the importance of the phylogenetic proximity. For example, researchers emphasized the role of the right IFG and the right orbitofrontal cortex (OFC), part of the PFC regions, in the human ability to correctly discriminate agonistic or affiliative contents in chimpanzee screams only (Belin, Fecteau, et al., 2008; Fritz et al., 2018). Nevertheless, Linnankoski and colleagues have shown the abilities of human adults and infants to recognize affective cues in macaque vocalizations using a categorization paradigm (Linnankoski et al., 1994). This last result points out the difference of complexity between the discrimination and categorization tasks in humans, even if the affective recognition is related to primate vocalizations. Overall, more controlled investigations in this domain are thus needed (Gruber & Grandjean, 2017).

Considering the paucity of neuroscientific studies adopting a comparative approach, the aim of the present study was to test the following questions using fNIRS: how are the human IFG and PFC regions involved in the explicit decoding of emotions contained in primate vocalizations? Is phylogenetic proximity a key for a better understanding of such processes? How does task complexity modulate the brain and behavioural responses across species and affect? To do so, we investigated human affective recognition processing in human and other primate vocalizations using cerebral and behavioural data. The participants performed categorization and discrimination tasks on affective contents (agonistic versus affiliative) in human, great apes (chimpanzee, bonobo) and monkey (rhesus macaque) vocalizations while their brain activity was recorded using fNIRS. We predicted that: i) according to the cognitive complexity hypothesis, the categorization task should involve more activations in the IFG and PFC than discrimination; ii) if a phylogenetic effect was at play, IFG and PFC would be modulated differently across human, great apes and monkey vocalizations; and iii) if frontal regions are necessary to cross-taxa recognition of affects, neural activity in the IFG and PFC should be related to the participants’ performances.

## Material & Methods

### Participants

Thirty healthy volunteers (12 males; mean age 25.06 years, SD = 5.09, age range 20-36) took part in the experiment. The participants reported normal hearing abilities and normal or corrected-to-normal vision. No participant presented a neurological or psychiatric history, or a hearing impairment. All participants gave informed and written consent for their participation in accordance with the ethical and data security guidelines of the University of Geneva. The study was approved by the Ethics Cantonal Commission for Research of the Canton of Geneva, Switzerland (CCER).

### Vocalizations

Ninety-six vocalizations of four primate species (human, chimpanzee, bonobo, rhesus macaque) in agonistic and affiliative contexts were used as stimuli. The human voices obtained from the Montreal Affective Voices (Belin, Fillion-Bilodeau, et al., 2008) were denoted as expressing a happy, angry or fearful affect (non-linguistic affective bursts) produced by two male and two female actors.

Vocalizations in corresponding contexts were selected for chimpanzee, bonobo and rhesus macaque species under the form of affiliative calls (food grunts), threatening calls (aggressor in agonistic context) and distress calls (victim in agonistic context). For each species, 24 stimuli were selected containing single calls or call sequences produced by 6 to 8 different individuals in their social environment.

All vocal stimuli were standardized to 750 milliseconds using PRAAT (www.praat.org) but were not normalized in order to preserve the naturalness of the sounds (Ferdenzi et al., 2013).

### fNIRS acquisition

fNIRS data were acquired using the Octamon device (Artinis Medical Systems B.V., Elst, The Netherlands) at 10 Hz with 6 transmitters and 2 receivers (wavelengths of ±760 nm and ±850 nm) with an inter-distance probes at 3.5 cm. The headband holding the 8 channels was placed identically for all participants according to the 10-20 electroencephalogram (EEG) system (Jasper, 1958; Okamoto et al., 2004) by using the FPZ axis as landmark (see Figure 1). The probe locations into the Montreal Neurological Institute (MNI) space were estimated using the 3D coordinates extracted from 32 healthy participants (Vergotte et al., 2018). Hence, the channels 1, 2, 7 and 8 were located on IFG_tri_ and the channels 3, 4, 5 and 6 on the PFC.

**Figure 1:**
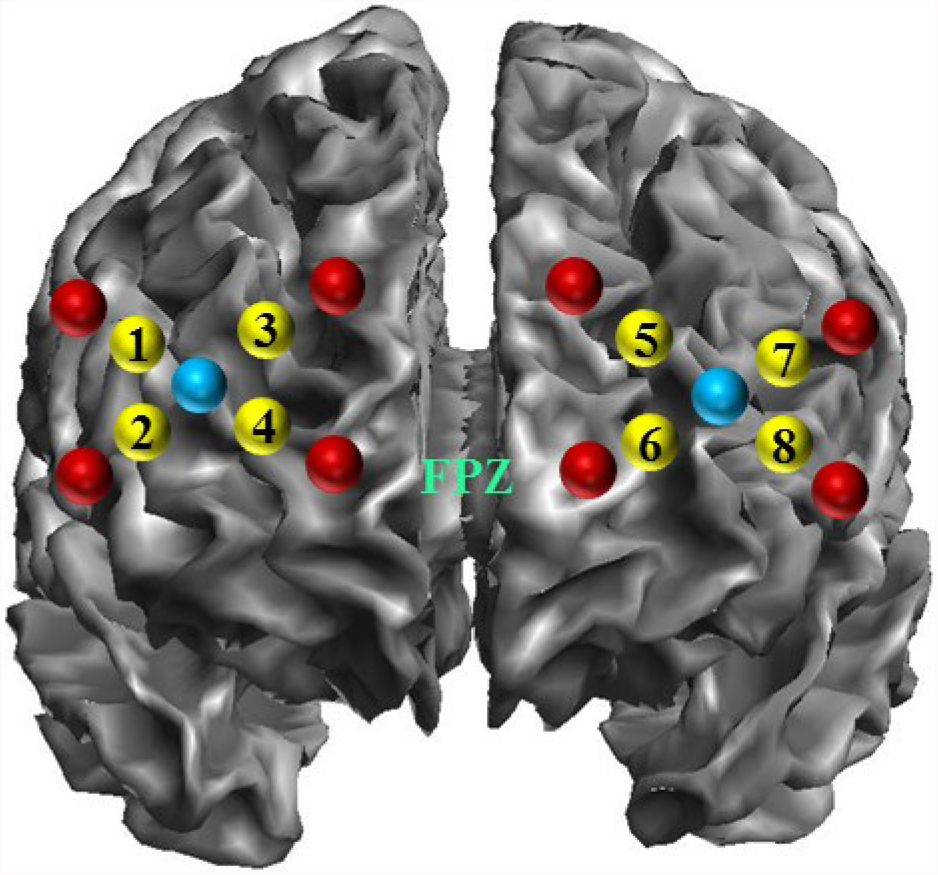
Probe locations into the MNI space by using SPM12 software implemented in MatLab R2018b (www.fil.ion.ucl.ac.uk/spm/). Red and blue dots indicate transmitters and receivers’ positions respectively. Yellow dots indicate the channel numbers.

### Experimental procedure

Seated comfortably in front of a computer, participants listened to the vocalizations played binaurally using Seinnheiser headphones at 70 dB SPL. Each of the 96 stimuli was repeated nine times across six separate blocks leading to 864 trials following a randomization process. The overall experiment was structured in various layers (Figure 2). Testing blocks were task-specific, with participants having to either perform a categorization task (A versus B) or a discrimination task (A versus non-A) in a single block. Participants completed three categorization blocks and three discrimination blocks, resulting in six blocks in total. Each block was made of 12 mini-blocks, each separated by a break of 10 seconds. These mini-blocks comprised one unique mini-block per species (human, chimpanzee, bonobo and rhesus macaque), each mini-block repeated 3 times. Within each mini-block were 12 trials, containing four vocalisations from all three affective contexts (affiliative/happy; threatening/anger; fear) produced by a single species. The blocks, mini-blocks and stimuli were pseudo-randomly assigned for each participant to avoid more than two consecutive blocks, mini-blocks and stimuli from the same category.

**Figure 2:**
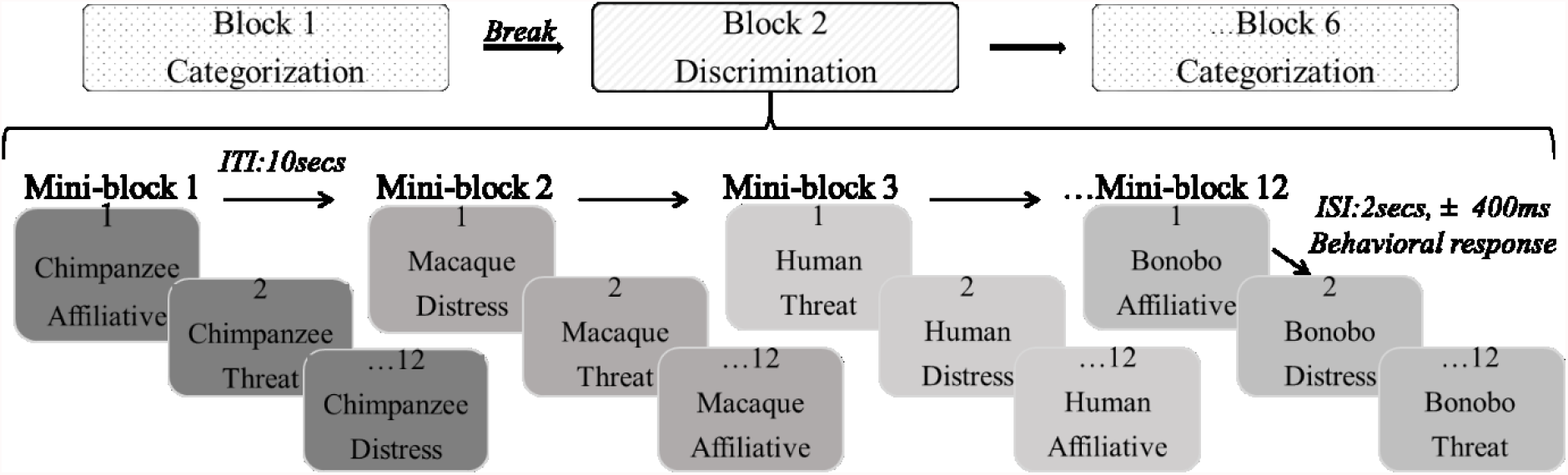
Structure of the experiment, with each of the six blocks made of 12 mini-blocks, which in turn comprised 12 individual trials.

At the beginning of each block, participants were instructed to identify the affective content of the vocalizations using a keyboard. For instance, the instructions for the categorization task could be “Affiliative – press M or Threatening – press Z or Distress – press space bar”. Similarly, the instructions for discrimination could be “Affiliative – press Z or other affect – press M”. The pressed keys were randomly assigned across blocks and participants. The participants had to press the key during the 2-second intervals (jittering of 400 ms) between each stimulus. If the participant did not respond during this interval, the next stimulus followed automatically.

### Statistical analysis

#### Behavioural data

Raw behavioural data from all participants were analysed using Generalized Linear Mixed Model (GLMM) fitted by Restricted Maximum Likelihood (REML) on R.studio (Team, 2020) with the “bobyqa” function (optimization by quadratic approximation with a set maximum of 1’000’000 iterations) and the link “logit” for a standard logistic distribution or errors and a binomial error distribution (correct answer – 1 or not – 0) of the package Lme4 (Bates et al., 2015). The following three factors and their interactions were included: Species (human, chimpanzee, bonobo, and rhesus macaque), Tasks (categorization - CAT and discrimination - DIS), and Affects (affiliative, threat, and distress). Participant IDs and order of the blocks were used as random factors. In order to test our hypotheses regarding the phylogenetic distance and the task complexity on participants’ performances we compared, using contrasts, the differences between Species and Affects within the categorization and the discrimination tasks. These contrasts were corrected with Bonferroni correction (P_corrected_ = .05/number of tests = .05/24=.002). Similarly, the participants’ reaction time (correct answers only) were analysed using a GLMM with a Gaussian distribution with the same contrasts and analysis as for accuracy. The present paper focusing on the investigation of recognition mechanisms, not attentional processes, results for reaction times are reported in supplementary material.

#### fNIRS data

Ten participants out of 30 were excluded from the dataset due to poor signal quality (large number of artefacts after filtering) or missing fNIRS data. A total of 20 participants were thus analysed in this study, in line with previous power analyses in fMRI (Desmond & Glover, 2002) and research using fNIRS to assess emotional processing in frontal areas (for a review, see Bendall et al., 2016). We performed on all channels the first level analysis with MatLab 2018b (Mathwortks, Natick, MA) using the SPM_fNIRS toolbox (Tak, Uga, Flandin, Dan, & Penny, 2016; https://www.nitrc.org/projects/spm_fnirs/) and homemade scripts. Haemoglobin conversion and temporal pre-processing of O_2_Hb was made using the following procedure:

1. Haemoglobin concentration changes were calculated with the modified Beer-Lambert law (Delpy et al., 1988);
2. Motion artefacts were reduced using the movement artefact reduction algorithm (MARA - Scholkmann et al., 2010) based on moving standard deviation and spline interpolation;
3. Low frequency confound were reduced using a high-pass filter based on a discrete cosine transform set with a cut-off frequency of 1/64 Hz (Friston et al., 2000);
4. Physiological and high frequency noise such vasomotion or heart beats usually found in extra-cerebral blood flow were removed using a low-pass filter based on the hemodynamic response function (HRF - Friston et al., 2000).
5. O_2_Hb concentration changes were averaged between 4 and 12 seconds post stimulus onset on each trial to include the maximum peak amplitude of the HRF observed across participants. As for fMRI imaging, this method of analysis taking into account the slow hemodynamic time course of brain activity is in line with previous literature using auditory stimuli in fNIRS (e.g. Lloyd-Fox et al., 2014).

The second level analysis was performed on R. studio using GLMM fitted by REML with the factors: Species (human, chimpanzee, bonobo, rhesus macaque), Tasks (categorization versus discrimination), Affects (affiliative, threatening, distressful), as well as their interactions as fixed factors, and participant IDs and block orders as random factors for the right and left IFG_tri_ and PFC.

#### Interaction between participants’ performance and brain Oxyhemoglobin (O_2_Hb) changes

To test whether the IFG_tri_ and PFC activations facilitated the participants’ affective recognition, we used fNIRS data as continuous predictors in GLMM analysis performed on R. studio for accuracy. To perform this statistical interaction, we only used accuracy from the twenty participants included in fNIRS analyses. The GLMM fitted by REML included Species (human, chimpanzee, bonobo and rhesus macaque), Tasks (discrimination and categorization), Affects (threat, distress and affiliative), as fixed factors, fNIRS data from the right and left IFG_tri_ and PFC as continuous predictors, and participant IDs as a random factor. To assess the variance explained by the phylogeny as well within the frontal activation, we tested all slopes with the following contrast: human *vs* [great apes (chimpanzee and bonobo)] *vs* rhesus macaque.

## Results

### Accuracy

We investigated how the perceptual decision-making complexity influenced the ability of human participants to recognize affective contents in phylogenetically close or distant primate species (see Figure 3).

**Figure 3:**
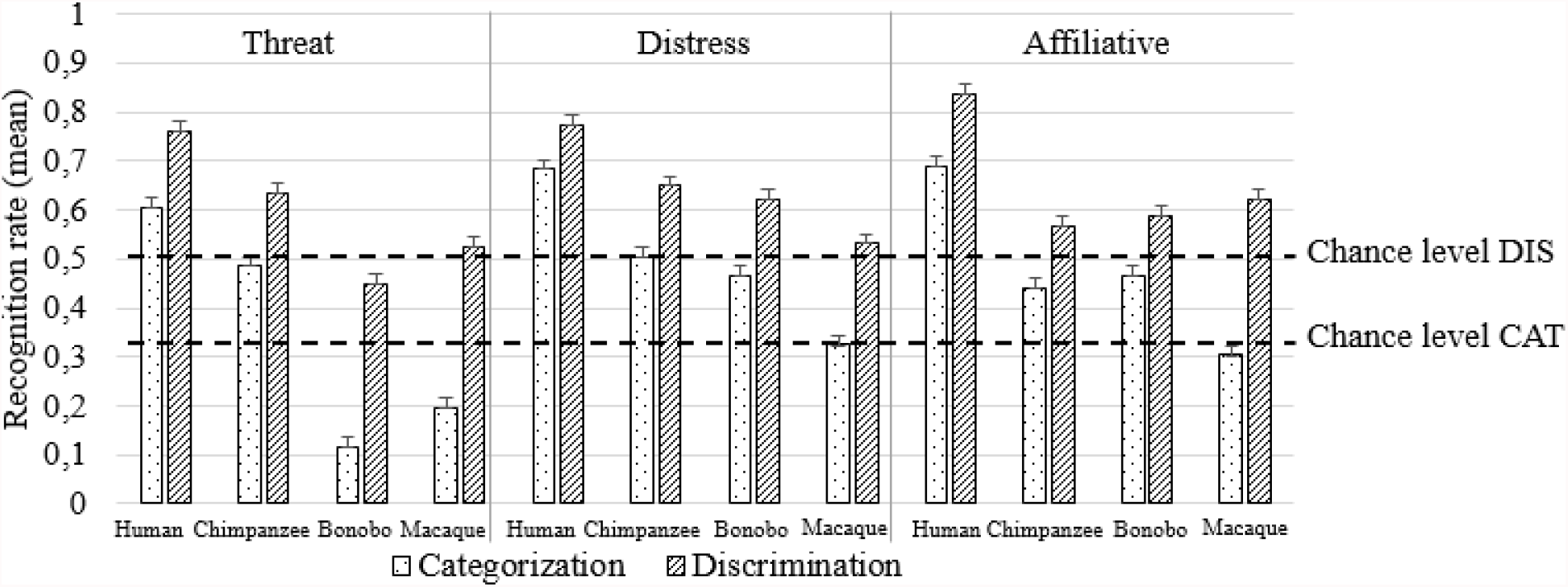
Mean and SE of human recognition of primate affective vocalizations for categorization (CAT) and discrimination (DIS) tasks and the different kinds of affective vocalizations. All contrasts were significant within each condition after Bonferroni correction with P_corrected_ = .05/24=.002, excluding the following contrasts: chimpanzee *vs* macaque and bonobo *vs* macaque for affiliative cues and bonobo *vs* macaque for threatening contents in discrimination task (see supplementary material Table 1).

Hence, participants were significantly above chance (>50% in discrimination; >33% in categorization) for most of the affective cues in great ape vocalizations (threatening bonobo calls excluded - see Table 1). Yet, they were unable to do so for threatening macaque calls in the discrimination task and all affective vocalizations expressed by this species in the categorisation one. Moreover, human participants were better at discriminating and then categorizing human voices (threat = DIS 76%; CAT 60%, distress = DIS 77%; CAT 68%, affiliative = DIS 83%; CAT 69%), chimpanzee distress (DIS 65%; CAT 50%) and threatening (DIS 63%; CAT 50%) vocalizations, followed by distress and affiliative calls expressed by bonobos (DIS 62%; CAT 46% for both) and macaques in the discrimination task (62%).

**Table 1:** Summary of the one sample t-test analyses against chance level. Recognition performance above chance (>33% categorisation and >50% discrimination) are written in bold. *** p < .001, * p < .05.

|  | Categorization |  |  | Discrimination |  |  |
| --- | --- | --- | --- | --- | --- | --- |
|  | Threat | Distress | Affiliative | Threat | Distress | Affiliative |
| Bonobo | -22.19*** | **8.95\*\*\*** | **8.96\*\*\*** | -3.44*** | **8.26\*\*\*** | **5.9\*\*\*** |
| Chimpanzee | **10.27\*\*\*** | **11.41\*\*\*** | **7.35\*\*\*** | **9.15\*\*\*** | **10.19\*\*\*** | **4.51\*\*\*** |
| Human | **18.47\*\*\*** | **25.07\*\*\*** | **25.67\*\*\*** | **19.92\*\*\*** | **21.33\*\*\*** | **29.51\*\*\*** |
| Macaque | -11.15*** | -0.41 | -1.93 | 1.53 | **2.02\*** | **8.13\*\*\*** |

### fNIRS data

A significant main effect was found for the factor Tasks in the right IFG_tri_ (χ^2^(1) = 14.27, p < .001)_;_ left IFG_tri_ (χ^2^(1) = 3.89, p < .05); right PFC (χ^2^(1) = 107.32, p < 0.001) and left PFC (χ^2^(1) = 90.83, p < .001) revealing more O2Hb concentration changes for the discrimination compared to the categorization task for all ROIs (see Figure 4). Note that none of the interactions with the factors Affects and Species reached significance.

**Figure 4:**
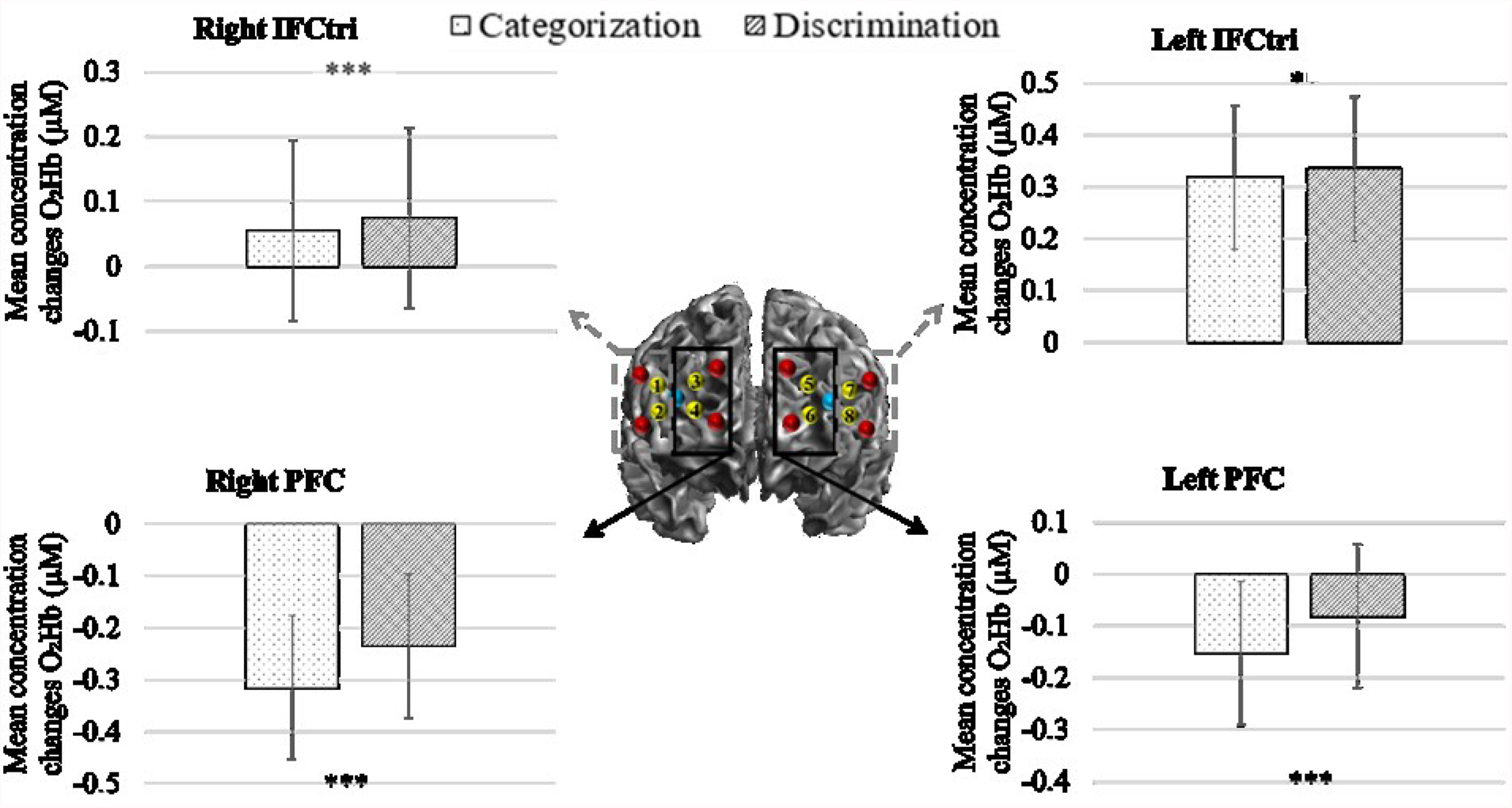
Mean and SE of concentration changes of O_2_Hb (μM) in right and left PFC and IFG_tri_ during the categorization and the discrimination tasks by human participants of primate affective vocalizations. *** p< .001, * p< .05.

### Interaction between participants’ performance and brain O_2_Hb changes

All factors (Tasks, Species and Affects) with the fNIRS data of the right and left IFG_tri_ and PFC as continuous predictors contributed to a significant three-way interaction (χ^2^(24) = 202,28 p <.001).

Within this model, we then assessed how the affective contents modulated IFG_tri_ and PFC activity across species vocalizations during the categorization or discrimination tasks. For this purpose, we investigated whether the participants’ accuracy and the related fNIRS data were positively, negatively or not correlated for each Species and ROIs within the Affects and Tasks factors using odd-ratio summarized in Table 2. In particular we tested whether phylogenetic proximity facilitated the recognition of Affect. We found for both the IFG_tri_ and PFC that contrasts between humans *vs* [great apes (chimpanzees and bonobos)] *vs* rhesus macaques within each Affect and Task were significant at p < .001 (see supplementary material Table 3). Note that because we found similar patterns of performances between PFC and IFG_tri_, for more clarity, we will only describe the results for IFG_tri_ here (see Figure 5). Results for PFC are reported in supplementary material Figure 3.

**Table 2:** Summary of the odds ratio and p-values testing the statistical significance and the direction of logistic regression slopes from the three-way interaction. The odds ratio quantifies the strength of the association between two factors. If the slope is significant and odds ratio < 1, factors are negatively correlated (written in bold); if the slope is significant and odds ratio > 1, factors are positively correlated (written in bold italic). ** p < .01, * p < .05.

|  | Categorization |  |  | Discrimination |  |  |
| --- | --- | --- | --- | --- | --- | --- |
|  | Threat | Distress | Affiliative | Threat | Distress | Affiliative |
| Bonobo | **0.84\*** | **0.88\*** | 1.06 | 0.99 | 1.1 | 1.06 |
| Chimpanzee | **0.78\*** | **0.69\*\*** | **0.86\*** | **1.28\*** | **1.44\*\*** | 0.93 |
| Human | 1.02 | 1.13 | 1.11 | 0.98 | 0.89 | 1.02 |
| Macaque | 1.07 | 0.94 | **0.85\*** | 0.93 | 0.9 | 1.05 |

**Figure 5:**
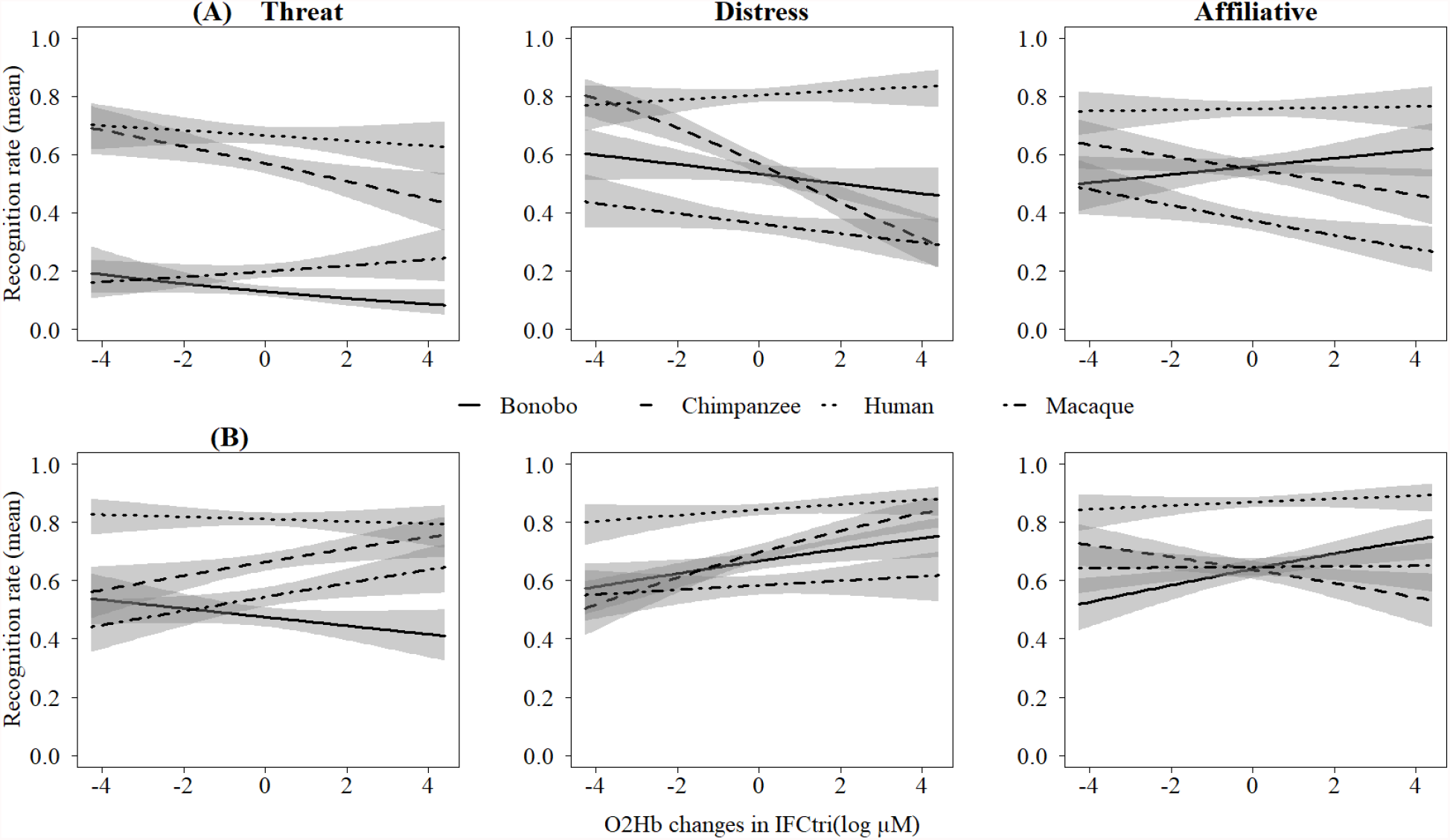
Interaction between participants’ accuracy and O_2_Hb concentration changes in IFG_tri_ within each affect and species for (A) categorization and (B) discrimination. Confidence interval at 0.95. Figures were made on R.studio using the package Visreg (Breheny & Burchett, 2017).

Hence, participants better discriminated agonistic (threat and distress) chimpanzee calls when the concentration changes of O_2_Hb increased in IFG_tri_ and PFC. At the opposite, during the categorization task, the correct identification of all types of chimpanzee calls as well as affiliative macaque and agonistic bonobo vocalizations were associated with a decrease of activity in frontal regions.

## Discussion

The present study emphasized the different levels of complexity in decision-making processes underlying the human recognition of affects in human and non-human primate vocalizations. In particular, we demonstrated that the left IFG_tri_ and the right PFC were strongly involved in the discrimination task compared to the categorization one.

Interestingly, and perhaps, contradictorily, we initially expected more activation in IFG_tri_ for the categorization task (unbiased choice) because of the existing literature on human affective voices (Dricu et al., 2017; Gruber et al., 2020). However, taking into account our behavioural results showing higher recognition performances in discrimination compared to categorization, more activity in IFG_tri_ appears to be required to enable participants to perform better during the discrimination of primate vocalizations. At the opposite, in line with the cognitive complexity hypothesis, analyses for PFC revealed a stronger deactivation in the categorization task. We could link these last findings to the changes in regional cerebral blood flow. Indeed, Matsukawa and collaborators showed that during the passive viewing of emotional videos, the activity of PFC decreased in correlation to the reduction of facial skin blood flow (Matsukawa et al., 2018). Interestingly, these authors suggested that PFC activity might elicit an autonomic reaction with a vasoconstriction or a vasodilatation of cutaneous vessels. In the same line, George and collaborators demonstrated a stronger decrease of activity in right PFC during the viewing of pleasant pictures, also relying on a reduction of the frontal blood flow (George et al., 1995). A possibility is thus to extend the results of these visual studies to a decrease of activity in PFC regions during affective auditory processing.

Overall, our results highlight the distinct roles of the IFG_tri_ and the PFC in evaluative judgment and decision task in affective primate calls recognition (see Schirmer & Kotz, 2006; Wagner & Watson, 2010 for humans).

Was human recognition influenced by the affects and/or the species that expressed the vocalizations? We did find an influence of these factors on behavioural responses and the interaction between participants’ performances and frontal activations. In fact, we demonstrated that the correct categorization of agonistic cues in bonobo and chimpanzee vocalizations elicited a significant decrease of activity in the IFG_tri_ and the PFC. These results might be related to an inhibition process enabling participants to reduce a high level of stress elicited by agonistic calls, i.e. automatic regulation. Frontal regions are indeed the most sensitive brain areas to stress exposure (Arnsten, 2009). Interestingly, a decrease of activation in frontal regions was also associated to better performance in the categorization task for affiliative chimpanzee and macaque vocalizations. On the contrary, in the discrimination task, agonistic chimpanzee screams were better identified when the level of activity in IFG_tri_ and PFC increased. These results highlight the involvement of distinct mechanisms between the categorization and discrimination tasks in cross-taxa recognition. For instance, possible inhibition processes elicited by agonistic cues would rely on a decrease of activations in frontal regions for the simple choice between A versus non-A; while in categorisation (unbiased choice), similar inhibition mechanisms would require an enhancement of activity in IFG_tri_ and PFC.

The general absence of interaction between frontal activations and behaviours for human voices might be explained by three different mechanisms. First, for humans, because affective voices in our modern human societies are everywhere (Belin, 2006), the correct recognition of affects may not necessary involve particular frontal activations due to the human expertise in human voice processing. Second, the involvement of IFG has often been demonstrated in the literature for the recognition of emotional voices contrasted with neutral ones (e.g. Frühholz et al., 2012; Frühholz & Grandjean, 2013; Gruber et al., 2020; Sander et al., 2005; Zhang et al., 2018). Yet, in our study, we did not include such stimuli, comparing cerebral activations across the affective contents. This difference in our experimental paradigm may have led to the absence of interaction between the hemodynamic response in the frontal regions and the emotional recognition in human voices. Third, encompassing three neuroanatomical and functional subparts: *pars triangularis, pars orbitalis* and *pars opercularis* (Cai & Leung, 2011), IFG_tri_ would possibly requires the recognition of infrequent vocalizations expressed by evolutionary close species to be modulated. Following this, the phylogenetic gap of 25-33 million between rhesus macaque and the *Hominidae* branch might explain the lack of result for this monkey species. Performances on the macaque calls categorization were poor, hence the frontal activations would not help to categorize them because human participants were, at least in this experiment, unable to categorize these calls. In contrast, participants were able to categorize most affects in great ape vocalizations, to the exception of threatening bonobo calls.

Yet, such reasoning does not apply to discrimination, where the low level of cognitive complexity involved may have allowed participants to discriminate more correctly affective vocalizations of all primates, including species with larger phylogenetic distances such as macaques. Strikingly, behavioural analyses revealed that human participants were able to discriminate most of the affective cues in all species vocalizations, once again to the exception of threatening bonobo calls. We might hypothesize that specific acoustic factors in bonobo calls triggered this effect: bonobo calls have indeed a higher fundamental frequency resulting from a shorter vocal length in comparison to chimpanzees. In this species, signalling physical strength using low frequencies (e.g. Briefer, 2012; Morton, 1982) is not a sexually selected trait (Grawunder et al., 2018). This reflects in their general behaviour, with bonobos being quite different from closely related chimpanzees and overall less aggression prone: they are occasional hunters, do not have strict territories and have a developed socio-sexuality, reducing the number of aggressive conflicts (Gruber & Clay, 2016).

To conclude, our findings demonstrate the interplay between cerebral and behavioural processes during the recognition by humans of affective cues in primate vocalizations. Decision-making complexity, phylogeny and behaviour seem four essential markers to consider for further studies on cross-taxa recognition. Overall, we demonstrated the difference of mechanisms between the categorization and discrimination of primate affective calls at both behavioural and cerebral levels. In particular, we showed various activations in the PFC and IFG_tri_ and their connection to the ability of humans to correctly identify affective cues in great apes and monkeys’ vocalizations. Furthermore, our results highlighted the importance of the phylogenetic proximity in affective recognition processes. Finally, to our knowledge, this study is the first to: i) distinguish categorization and discrimination processes in a neuroscientific experiment with a comparative perspective, and ii) to assess the link between cross-taxa affective recognition and frontal activations in a fNIRS paradigm. We hope these new findings will contribute to a better understanding of the evolutionary origins of emotional processing and decision-making origin in human, as well as advocate for the inclusion of a broader array of auditory stimuli.

## Acknowledgements

We thank Katie Slocombe very much for providing chimpanzee and macaque auditory stimuli as well as extensive comments on former versions of this preprint. We would like also to thanks Dr. Ben Meuleman for his useful support on statistical analyses. We thank the Swiss National Science foundation (SNSF) for supporting this interdisciplinary project (CR13I1_162720 / 1 – DG-TG), and the Swiss Center for Affective Sciences. ZC has received support from the ESRC-ORA (ES/S015612/1), the ERC Starting Grant (802979, and CD from the foundation Ernst and Lucie Schmidheiny. TG was additionally supported by a grant of the SNSF during the final re-writing of this article (grant PCEFP1_186832).

