## Supplementary material for "Categorization and discrimination of human and non-human primate affective vocalizations: investigation of the frontal cortex activity through fNIRS"

*Accuracy*

Table 1: Main contrasts for the participants' accuracy associated with degree of freedom, chi-squared tests and p-values for the interaction Species x Affect x Task. All p-values are corrected for multiple comparisons using Bonferroni correction with $P_{corrected} = 0.002$.

| **Contrasts** | **Dfs** | $\chi^2$ | **p-values** |
|---|---|---|---|
| **Categorization** | | | |
| **Threat** | | | $p < 0.001$ |
| **Human > Chimpanzee** | 1 | 27.844 | $p < 0.001$ |
| **Human > Bonobo** | 1 | 506.96 | $p < 0.001$ |
| **Human > Macaque** | 1 | 363.35 | $p < 0.001$ |
| **Chimpanzee > Macaque** | 1 | 210.18 | $p < 0.001$ |
| **Bonobo > Macaque** | 1 | 28.064 | $p < 0.001$ |
| **Distress** | | | |
| **Human > Chimpanzee** | 1 | 84.845 | $p < 0.001$ |
| **Human > Bonobo** | 1 | 135.8 | $p < 0.001$ |
| **Human > Macaque** | 1 | 319.29 | $p < 0.001$ |
| **Chimpanzee > Macaque** | 1 | 90.67 | $p < 0.001$ |
| **Bonobo > Macaque** | 1 | 50.304 | $p < 0.001$ |
| **Affiliative** | | | |
| **Human > Chimpanzee** | 1 | 106.2 | $p < 0.001$ |
| **Human > Bonobo** | 1 | 91.628 | $p < 0.001$ |
| **Human > Macaque** | 1 | 287.96 | $p < 0.001$ |
| **Chimpanzee > Macaque** | 1 | 51.584 | $p < 0.001$ |
| **Bonobo > Macaque** | 1 | 64.275 | $p < 0.001$ |
| **Discrimination** | | | |
| **Threat** | | | |
| **Human > Chimpanzee** | 1 | 42.652 | $p < 0.001$ |
| **Human > Bonobo** | 1 | 210.35 | $p < 0.001$ |
| **Human > Macaque** | 1 | 139.03 | $p < 0.001$ |
| **Chimpanzee > Macaque** | 1 | 31.205 | $p < 0.001$ |
| **Bonobo > Macaque** | 1 | 9.4646 | $p < 0.01$ |
| **Distress** | | | |
| **Human > Chimpanzee** | 1 | 35.233 | $p < 0.001$ |
| **Human > Bonobo** | 1 | 64.306 | $p < 0.001$ |
| **Human > Macaque** | 1 | 133.29 | $p < 0.001$ |
| **Chimpanzee > Macaque** | 1 | 34.8 | $p < 0.001$ |
| **Bonobo > Macaque** | 1 | 14.751 | $p < 0.001$ |
| **Affiliative** | | | |
| **Human > Chimpanzee** | 1 | 152.76 | $p < 0.001$ |

| | | | |
|---|---|---|---|
| **Human > Bonobo** | 1 | 143.37 | $p < 0.001$ |
| **Human > Macaque** | 1 | 104.94 | $p < 0.001$ |
| **Chimpanzee > Macaque** | 1 | 5.7725 | $p = 1$ |
| **Bonobo > Macaque** | 1 | 3.6636 | $p < 0.1$ |

*Reaction time*

Including only the correct answers of the participants, we firstly tested the difference between the categorization and discrimination tasks of affects in human voices and primate vocalizations. GLMM analysis revealed a reaction time less long in affective discrimination compared to categorization for human ($\chi^2(1) = 176.61$, $p < 0.001$); chimpanzee ($\chi^2(1) = 140.7$, $p < 0.001$); bonobo ($\chi^2(1) = 49.075$, $p < 0.001$) and rhesus macaque ($\chi^2(1) = 152.53$, $p < 0.001$) vocalizations.

For the subsequent contrasts testing the phylogenetic hypothesis, we found that participants were faster to categorize and discriminate human voices compared to chimpanzee (cat: $\chi^2(1) = 334.16$, $p < 0.001$, dis: $\chi^2(1) = 393.95$, $p < 0.001$); bonobo (cat: $\chi^2(1) = 184.11$, $p < 0.001$, dis: $\chi^2(1) = 448.5$, $p < 0.001$) and macaque (cat: $\chi^2(1) = 276.86$, $p < 0.001$, dis: $\chi^2(1) = 315.03$, $p < 0.001$) affective vocalizations. Further analyses also revealed better performance in discriminating affective contents in bonobo vocalizations compared to rhesus macaque calls with the contrast bonobo *vs* rhesus macaque ($\chi^2(1) = 10.811$, $p = 0.001$).

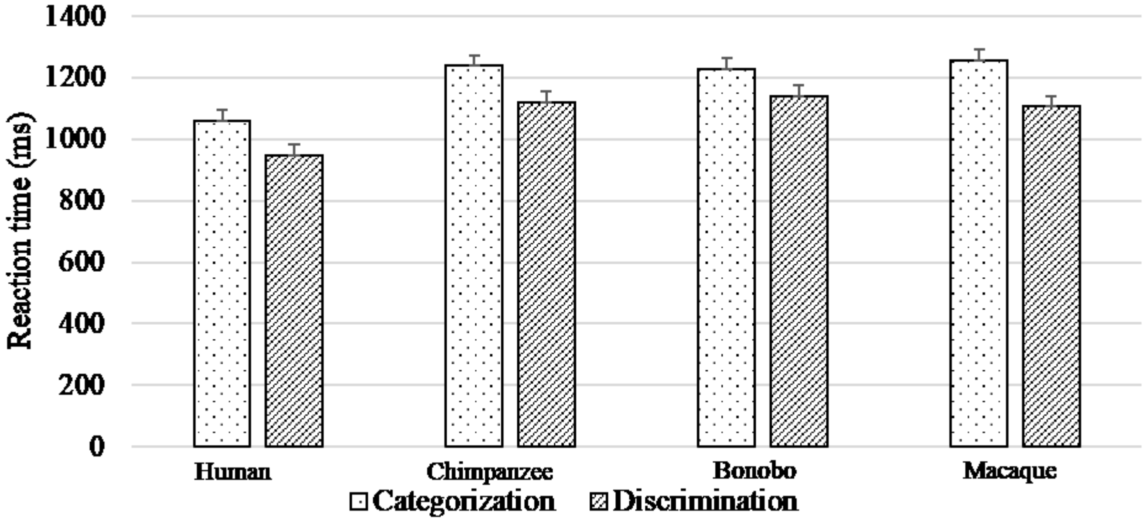

Figure 1: Means and standard errors of human reaction time (milliseconds) of primate affective vocalizations for categorization and discrimination tasks.

Finally, as for participants' accuracy we investigated how the different affects, species and tasks influenced the reaction time of human recognition.

Therefore, participants needed less time to discriminate than categorize affective contents in all species vocalizations. Thus, participants were faster at recognizing affective cues in human voices (cat = 1080ms, dis = 950ms) than chimpanzee (cat = 1210ms, dis = 1100ms), bonobo (cat = 1210ms, dis =980 ms) or macaque (cat = 1230ms, dis = 930ms) vocalizations. Statistics revealed difference in both tasks depending on species and affective cues. Participants were faster to discriminate (in this order) human voices (threat = 950ms, distress = 1000ms, affiliative = 900ms), affiliative macaque screams (1090ms), distressful bonobo calls (1100ms) and threatening chimpanzee vocalizations (1110ms). Considering the categorization task, similar findings were found with some differences only for chimpanzee and bonobo vocalizations.

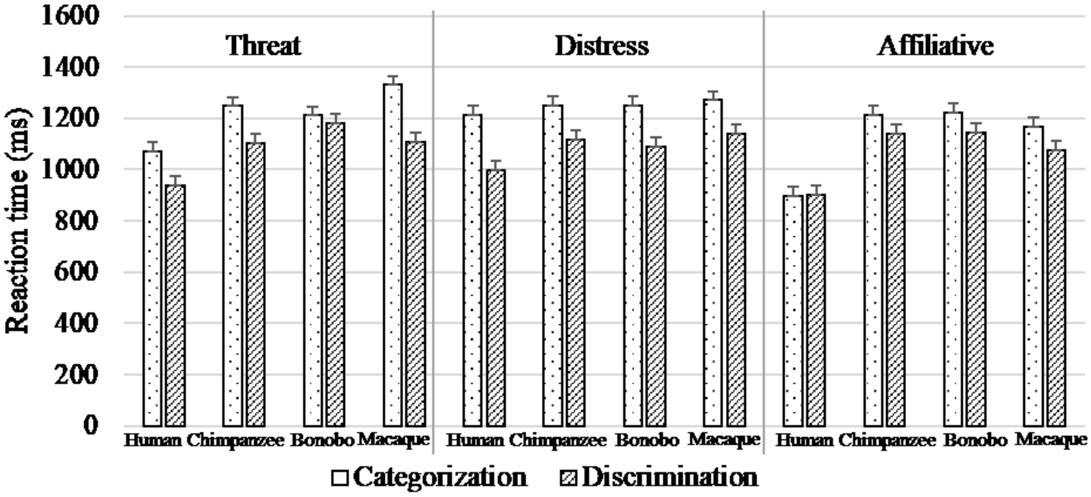

Figure 2: Means and standard errors of human reaction time (milliseconds) of primate affective vocalizations for categorization and discrimination tasks and the different kinds of affective vocalizations. All contrasts were significant within each condition after Bonferroni correction with $P_{corrected} = 0.002$, excluding the following contrasts: chimpanzee *vs* macaque and bonobo *vs* macaque for distressful cues in discrimination task. Regarding the categorization task, human *vs* chimpanzee, human *vs* bonobo, chimpanzee *vs* macaque and bonobo *vs* macaque for distressful cues and finally chimpanzee *vs* macaque and bonobo *vs* macaque for affiliative contents.

Table 2: Main contrasts for participants' accuracy associated with degree of freedom, chi-squared tests and p-values for the interaction Species x Affect x Task. All p-values are corrected for multiple comparisons using Bonferroni correction with $P_{corrected} = 0.002$.

| **Contrasts** | **Dfs** | $\chi^2$ | **p-values** |
|---|---|---|---|
| **Categorization** | | | |
| **Threat** | | | |
| **Human > Chimpanzee** | 1 | 104.45 | $p < 0.001$ |
| **Human > Bonobo** | 1 | 23.448 | $p < 0.001$ |
| **Human > Macaque** | 1 | 124.26 | $p < 0.001$ |
| **Chimpanzee > Macaque** | 1 | 12.164 | $p < 0.001$ |
| **Bonobo > Macaque** | 1 | 12.116 | $p < 0.001$ |
| **Distress** | | | |
| **Human > Chimpanzee** | 1 | 5.6393 | $p < 0.05$ |
| **Human > Bonobo** | 1 | 5.1341 | $p < 0.05$ |
| **Human > Macaque** | 1 | 9.717 | $p < 0.001$ |
| **Chimpanzee > Macaque** | 1 | 0.9859 | $p = 1$ |

| | | | |
|---|---|---|---|
| **Bonobo > Macaque** | 1 | 1.0431 | p < 1 |
| **Affiliative** | | | |
| **Human > Chimpanzee** | 1 | 350.83 | p < 0.001 |
| **Human > Bonobo** | 1 | 378.44 | p < 0.001 |
| **Human > Macaque** | 1 | 200 | p < 0.001 |
| **Chimpanzee > Macaque** | 1 | 5.0866 | p < 0.05 |
| **Bonobo > Macaque** | 1 | 6.7794 | p < 0.01 |
| **Discrimination** | | | |
| **Threat** | | | |
| **Human > Chimpanzee** | 1 | 118.3 | p < 0.001 |
| **Human > Bonobo** | 1 | 210.46 | p < 0.001 |
| **Human > Macaque** | 1 | 110.98 | p < 0.001 |
| **Chimpanzee > Macaque** | 1 | 0.0574 | p = 1 |
| **Bonobo > Macaque** | 1 | 16.954 | p < 0.001 |
| **Distress** | | | |
| **Human > Chimpanzee** | 1 | 63.03 | p < 0.001 |
| **Human > Bonobo** | 1 | 36.361 | p < 0.001 |
| **Human > Macaque** | 1 | 78.219 | p < 0.001 |
| **Chimpanzee > Macaque** | 1 | 1.6618 | p = 1 |
| **Bonobo > Macaque** | 1 | 8.6604 | p < 0.01 |
| **Affiliative** | | | |
| **Human > Chimpanzee** | 1 | 239.25 | p < 0.001 |
| **Human > Bonobo** | 1 | 254.39 | p < 0.001 |
| **Human > Macaque** | 1 | 130.74 | p < 0.001 |
| **Chimpanzee > Macaque** | 1 | 16.462 | p < 0.001 |
| **Bonobo > Macaque** | 1 | 19.252 | p < 0.001 |

*Accuracy * fNIRS data*

Table 3: Summary of contrasts testing phylogenetic differences with humans *vs* [great apes (chimpanzees and bonobos)] *vs* rhesus macaques between the slopes for the interaction Species x Affects x Tasks with fNIRS data for the right and left $IFC_{tri}$ and PFC as continuous predictors. All p-values are corrected for multiple comparisons using Bonferroni correction with $P_{corrected}$ =0.002.

| **$IFC_{tri}$** | | | |
|---|---|---|---|
| | Threat | Distress | Affiliative |
| Categorization | $\chi^2(1)$ = 609.25, p < 0.001 | $\chi^2(1)$ = 795.73, p < 0.001 | $\chi^2(1)$ = 1170.5, p < 0.001 |
| Discrimination | $\chi^2(1)$ = 456.57, p < 0.001 | $\chi^2(1)$ = 386.25, p < 0.001 | $\chi^2(1)$ = 499.69, p < 0.001 |
| **PFC** | | | |
| Categorization | $\chi^2(1)$ = 609.53, p < 0.001 | $\chi^2(1)$ = 797.88, p < 0.001 | $\chi^2(1)$ = 1166.4, p < 0.001 |

| | | | |
|---|---|---|---|
| Discrimination | $\chi^2(1) = 454.9, p < 0.001$ | $\chi^2(1) = 384.98, p < 0.001$ | $\chi^2(1) = 501.36, p < 0.001$ |

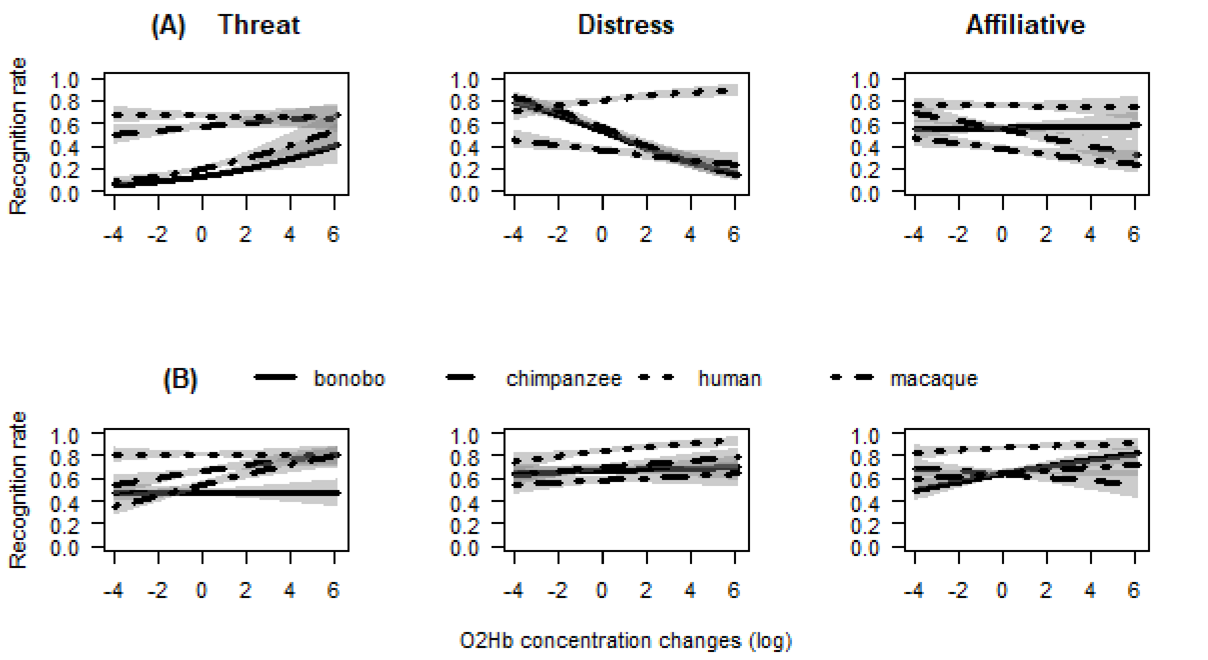

Figure 4: Interaction between participants' accuracy and concentration changes of $O_2Hb$ within Species and Affects for PFC in (A) categorization and (B) discrimination. Confidence interval at 0.95.
